# White matter correlates of hemi-face dominance in happy and sad expression

**DOI:** 10.1101/232926

**Authors:** Stefano Ioannucci, Nathalie George, Patrick Friedrich, Leonardo Cerliani, Michel Thiebaut de Schotten

## Abstract

The neural underpinnings of human emotional expression are thought to be unevenly distributed among the two brain hemispheres. However, little is known on the anatomy supporting this claim, particularly in the cerebral white matter. Here, we explored the relationship between hemi-face dominance in emotional expression and cerebral white matter asymmetries in 33 healthy participants. Measures of emotional expression were derived from pictures of the participant’s faces in a ‘happy smiling’ and a ‘sad frowning’ conditions. Chimeric faces were constructed by mirroring right and left hemi-faces, as done in previous studies, resulting in a left mirrored and right mirrored chimeric face per picture. To gain measures of hemi-face dominance per participant, a jury of 20 additional participants rated which chimeric face shows higher intensity of emotional expressivity, by marking a 155mm line between the two versions. Measures of the asymmetry of the uncinate, the cingulum and the three branches of superior longitudinal fasciculi were derived from diffusion weighted imaging tractography dissections. Group effect analyses indicated that the degree of asymmetry in emotional expression was not as prominent as reported in the literature and showed a large inter-individual variability. The degree of asymmetry in emotional expression was, however, significantly associated with the asymmetries in connective properties of the fronto-temporal and fronto-parietal tracts, specifically the uncinate fasciculus and the first branch of the superior longitudinal fasciculus. Therefore, this result raises novel hypotheses on the relationship of specific white matter tracts and emotional expression, especially their role in mood disorders.

**Ethical statement:**

- None of the authors have a conflict of interest
- Data collection from human participants was approved by the Comité de Protection des Personnes “CPP Ile de France V”
- All participants provided written informed consent
- Funding from ERC (grant agreement No. 818521) and “Agence Nationale de la Recherche” [grants numbers ANR-13-JSV4-0001-01 an ANR-10-IAIHU-06]

---

Over a century and a half ago, scientific research started to gather proof for the asymmetrical organization of the brain. Since the acknowledgement of Broca’s area (Broca 1861), the left hemisphere has been recognized as predominant in the language domain, whilst the right hemisphere has been linked to space, emotions, and their processing (Babinski 1914, Mills 1912, Sperry 1974).

Focusing on the emotion lateralization, a plethora of converging evidence has been gathered from cognitive, neuroanatomical, neuropsychological, clinical and behavioural studies, with substantial methodological diversity in paradigms, measures, populations, and emotional aspect investigated (e.g. perception, recognition, production or physiological response; for reviews please see Borod 1993, Demaree et al. 2005, Silberman and Weingartner 1986, Gainotti 2019).

Consequently, a variety of theoretical approaches have been developed to frame these findings, such as the *Right Predominance* hypothesis, the *Valence & Approach/Withdrawal* model (Demaree et al 2005, Silberman and Weingartner 1986) and the *Homeostatic Theory of Emotion* (Craig, 2005). This has resulted in a further increase in the array of experimental designs and research targets applied by the investigators. Hence, to this day, the concept of hemispheric dominance in emotion is still debated and regarded as antiquate by some cognitive scientists.

However, primates do express emotion asymmetrically (Bradshaw and Rogers 1993, Hellige 2001), indicating a phylogenetically old phenomenon. For instance, monkey chimeric facial stimuli (i.e. stimuli composed by mirroring the two halves of the face) reveal that they initiate their emotional expression earlier and end them later on the left side of their face (Fernández-Carriba et al. 2002, Hauser 1993). Further, a significant preference for the left hemi-spatial field even exists when chimpanzees—our closest primate ancestor—have to discriminate chimeric human smiling faces (Morris and Hopkins 1993). Therefore, given these premises, one might expect asymmetrical processing of emotional expression in humans.

In line with this expectation, studies in humans supporting facial asymmetries during emotional expression have accumulated over the years. Sackheim et al (1978) employed chimeric facial stimuli expressing 6 different emotions to be evaluated by 86 volunteers and demonstrated that the chimeric left/left hemi-faces were reported to be significantly more emotionally expressive. A comprehensive review on the existing literature by Borod et al (1997) concluded that the left hemiface is more involved than the right hemi-face in the expression of human facial emotion, implicating the right cerebral hemisphere as dominant in such a process. However, left and right hemi-faces emotional expressivity were always, to our knowledge, evaluated in isolation rather than in competition, thus limiting the conclusions since it did not favour a direct, paired comparison between left and right hemi-faces. In light of this potential caveat, a new paradigm comparing directly left and right hemi-faces is crucial to validate previous claims of right hemispheric dominance for the expression of emotions and to explore its relationship with brain anatomy.

The striking similarity in results regardless of species is suggestive of a common phylogenetic circuitry for emotional expression. It is widely accepted that emotion and related behaviour emerge from the joint activities of a cluster of brain regions known as the limbic system (Catani et al. 2013). Since its earliest conceptualizations, this system has been considered as a complex network of regions that act as an intermediary between a primitive, subcortical, brain and a more evolved, cortical one (MacLean 1952, Yakovlev 1948). Among the subcortical structures, within the inner anterior temporal lobe lies the amygdala, whose role in emotions has been largely documented (Phelphs 2006, Phelphs and LeDoux 2005). It is thought to be part of a functional subdivision of the limbic system, the temporo-amygdala-orbitofrontal network, connected by the uncinate fasciculus, which is involved in the integration of visceral and emotional states with cognition and behaviour (Mesulam 2000). Alterations in the aforementioned network have been linked to patients with clinical conditions involving emotional and mood disturbances, such as Fronto-Temporal Dementia (FTD) and psychopathy, among others (Catani et al. 2013). However, abnormalities in white matter pathways are found frequently not only within (i.e. uncinate fasciculus) but also outside (i.e. superior longitudinal fasciculus) the temporo-amygdala-orbitofrontal circuit in patients suffering psychopathy (Craig, 2009) or behavioural variant FTD (Agosta et al. 2011, Matsuo et al. 2008, Whitwell et al. 2010), whose deficits mainly affect emotional and social processing (Piguet et al. 2011). A second functional subdivision of the limbic system constitute parts of the default-mode network (DMN), a group of medial and lateral regions whose activity increases during passive thought not focused on a particular task (Raichle and Snyder 2007, Raichle et al. 2001) and has been associated with discrete emotions (Satpute and Lindquist, 2019). This network is connected by the cingulum and some part of the superior longitudinal fasciculus (SLF; Alves et al. 2019). Additionally, a meta-analysis by Jenkins et al (2016), found disruptions in the SLF as the most common white matter alteration in patients suffering from psychiatric emotional conditions such as bipolar disorder, social anxiety disorder, major depressive disorder, post-traumatic stress disorder and obsessive-compulsive disorder. Hence the superior longitudinal fasciculus, albeit frequently associated with spatial functions (Parlatini et al. 2017, Thiebaut de Schotten et al. 2011a) may have an emotional subcomponent.

Therefore, the aim of the present study was to contribute to the identification of anatomical biomarkers supporting hemi-face dominance in emotional expression in the human brain. We reconstructed *in vivo* estimates of white matter fibre pathways (Basser et al. 1994) that we statistically related to behavioural measures of emotional expression on chimeric facial stimuli. Particularly, we hypothesized that the hemispheric asymmetries of the uncinate, cingulum and superior longitudinal fasciculi relates to the hemispheric dominance of expression of emotions in the human brain.

## Methods

### Participants and Emotional expression task

The experiment was approved by the local ethics committee; all participants provided written informed consent in accordance to the Declaration of Helsinki. Participants also received an indemnity of 50 € for their participation.

A first group of 33 participants (later referred to as posers, mean age 27.6 ±5.03, 11 male) were included in this study. To obtain a measure of their emotional expression a picture of their faces in a ‘happy smiling’ and a ‘sad frowning’ condition was taken using a CANON PowerShot SD1790 IS 10.0 mega pixels. Participants were plainly instructed to pose in a happy and sad expression, as they would naturally do. These photos were edited to remove hair and the border of the face in order to maintain the characteristics that determine the saliency of emotions (e.g. eyes, mouth) and uniformed by normalization of grey levels reducing the saturation to minimum and adjusting the exposure parameters in the software *Preview.* According to previous attempts (Sackheim et al. 1978) pictures were split on the midline of the full face and chimeric stimuli were constructed by mirroring each half of the face. This resulted in two chimeric facial stimuli, one made of the left side and one made up of the right side (Figure 1a,b). This group also underwent diffusion MRI scans in order to reconstruct their white matter tracts. This material is available on demand to

**Figure 1:**
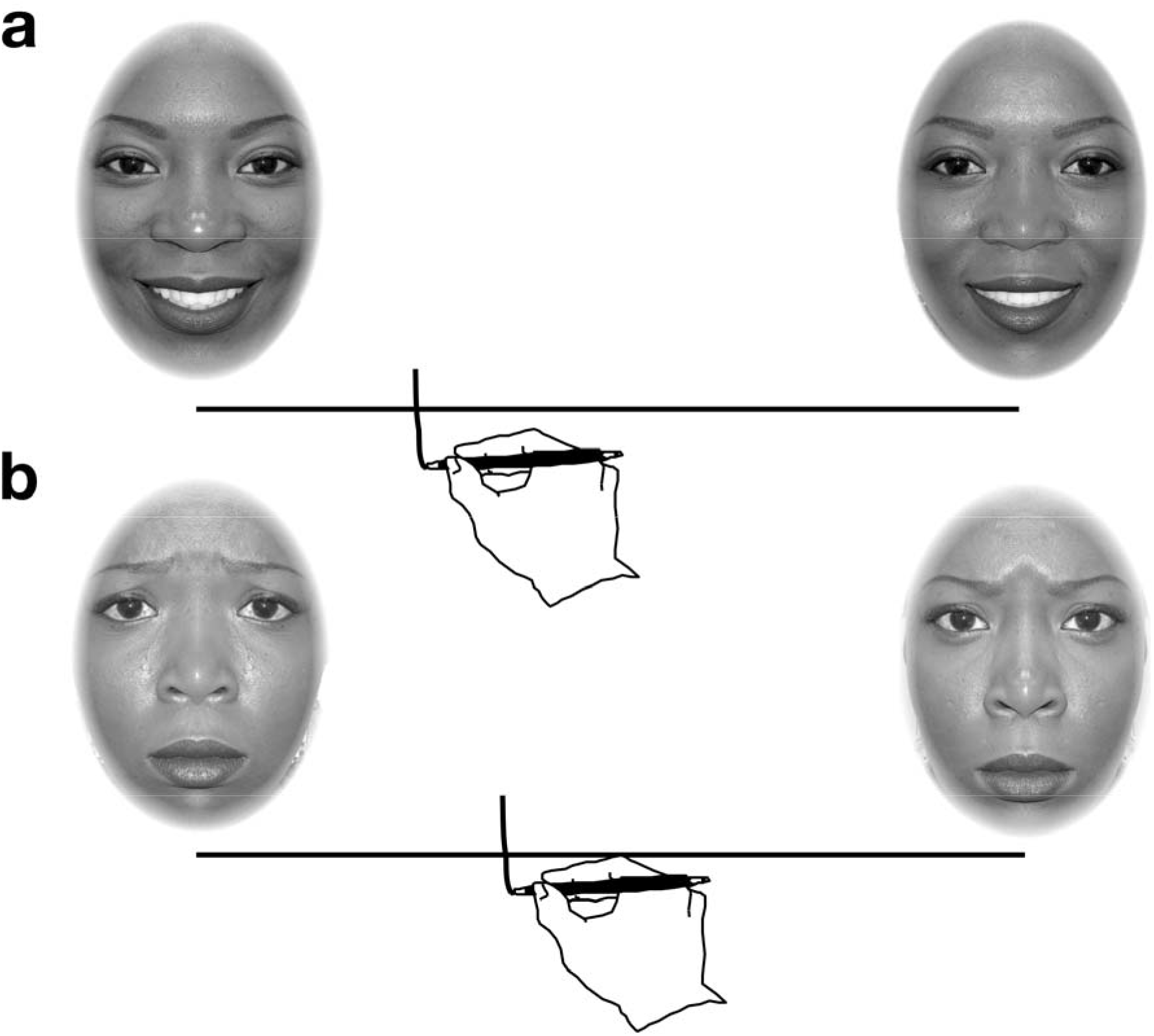
examples of chimeric facial stimuli asymmetry task for the happy (a) and sad (b) conditions. In this case, the stimuli placed on the left of the line are being valued as more emotionally expressive than the ones on the right, since the subject put his mark to the left of the centre of the line.

A second group of 20 participants (referred to as judges, mean age 25.65 ±4.6, 10 males) judged the set of chimeric facial stimuli from the group of posers. For this, they evaluated the Right/Right and Left/Left chimeric faces presented by pairs, at the extremities of a 155mm segment; they had to mark the segment according to the relative emotional intensity of the two stimuli, placing the mark closer to the stimuli that expressed the emotion more intensely, with the centre of the segment representing equal emotional intensity between the 2 stimuli. These stimuli were pseudo randomized. Stimuli were presented in a controlled procedure to the judges, half of the judges began scoring the faces from sad condition to happy condition while the other half did the opposite. These subgroups were further subdivided in half, so that some started with the stimuli being presented in a Left/Left – Right/Right hemi-face disposal, and reversed Right/Right – Left/Left disposal, in order to control for pseudoneglect effect (Jewell and McCourt, 2000). Therefore, the overall emotion scoring phase included 132 trials per judge (33 faces x 2 emotions x 2 disposals of these faces along the segment).

An asymmetry index was calculated for each face and each emotion expressed by the group of 33 posers, based on the averages of the deviations toward the chimeric stimuli that were rated higher in terms of emotional expressivity by the 20 judges. Reliability of this assessment was estimated using a correlation between a first and the second set of 10 judges.

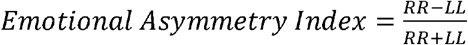

### RR and LL Represent the marks towards the stimuli as measured by distance from the middle of the segment

Therefore, a negative score indicated a tendency for left hemi-face dominance in emotional expression rated as more emotionally intense, while a positive score pointed to a right hemi-face dominance.

### Magnetic resonance imaging

Magnetic resonance imaging was acquired in the first group of 33 participants.

A total of 81 near-axial slices were acquired on a Siemens 3 Tesla Prisma system equipped with a 64-channel head coil. We used a fully optimised acquisition sequence for the tractography of diffusion-weighted imaging (DWI), which provided isotropic (1.7 × 1.7 × 1.7 mm) resolution and coverage of the whole head with a posterior-anterior phase of acquisition with an echo time (TE) = 75 msec. We used a repetition time (TR) equivalent to 3500ms. At each slice location, 6 images were acquired with no diffusion gradient applied (b-value of 0 sec mm-2). Additionally, 60 diffusion-weighted images were acquired, in which gradient directions were uniformly distributed on the hemisphere with electrostatic repulsion. The diffusion weighting was equal to a b-value of 2000 s/mm^-2^. This sequence was fully repeated with reversed phase-encode blips. This provides us with two datasets with distortions going in opposite directions. From these pairs the susceptibility-induced off-resonance field was estimated using a method similar to that described in (Andersson et al. 2003) and corrected on the whole diffusion weighted dataset using the tool TOPUP and EDDY as implemented in FSL (Smith et al. 2004).

Spherical deconvolution was chosen to estimate multiple orientations in voxels containing different populations of crossing fibres (Alexander 2005, Anderson 2005, Tournier et al. 2004). The damped version of the Richardson-Lucy algorithm for spherical deconvolution (Dell’Acqua et al. 2010) was calculated using Startrack (https://www.mr-startrack.com). Algorithm parameters were chosen as previously described (Dell’Acqua et al. 2012). A fixed-fibre response corresponding to a shape factor of α = 1.5 × 10–3 mm^2^/s was chosen (Dell’Acqua et al. 2012). Fiber orientation estimates were obtained by selecting the orientation corresponding to the peaks (local maxima) of the fiber orientation distribution (FOD) profiles. To exclude spurious local maxima, we applied both an absolute and a relative threshold on the FOD amplitude. A first “absolute” threshold was used to exclude intrinsically small local maxima due to noise or isotropic tissue. This threshold was set to 3 times the mean amplitude of a spherical FOD obtained from a grey matter isotropic voxel (and therefore also higher than an isotropic voxel in the cerebro-spinal fluid).

A second “relative” threshold of 10% of the maximum amplitude of the FOD was applied to remove maximum amplitude of the FOD was applied to remove the remaining local maxima with values greater than the absolute threshold (Dell’Acqua et al. 2010). Whole brain tractography was performed selecting every brain voxel with at least one fibre orientation as a seed voxel. From these voxels, and for each fibre orientation, streamlines were propagated using Euler integration with a step size of 1 mm (as described in (Dell’Acqua et al. 2012)). When entering a region with crossing white matter bundles, the algorithm followed the orientation vector of least curvature (Schmahmann et al. 2007). Streamlines were halted when a voxel without fibre orientation was reached or when the curvature between two steps exceeded a threshold of 60°. Spherical deconvolution, fibre orientation vector estimations and tractography were also implemented in Startrack.

### Tractography dissections

Virtual dissections were made of the bilateral tracts on which we had a priori hypotheses, that is, the uncinate fasciculus, the three branches of the superior longitudinal fasciculi (SLF I, II and III) and the cingulum (**Figure 2 a,b,c**).

**Figure 2:**
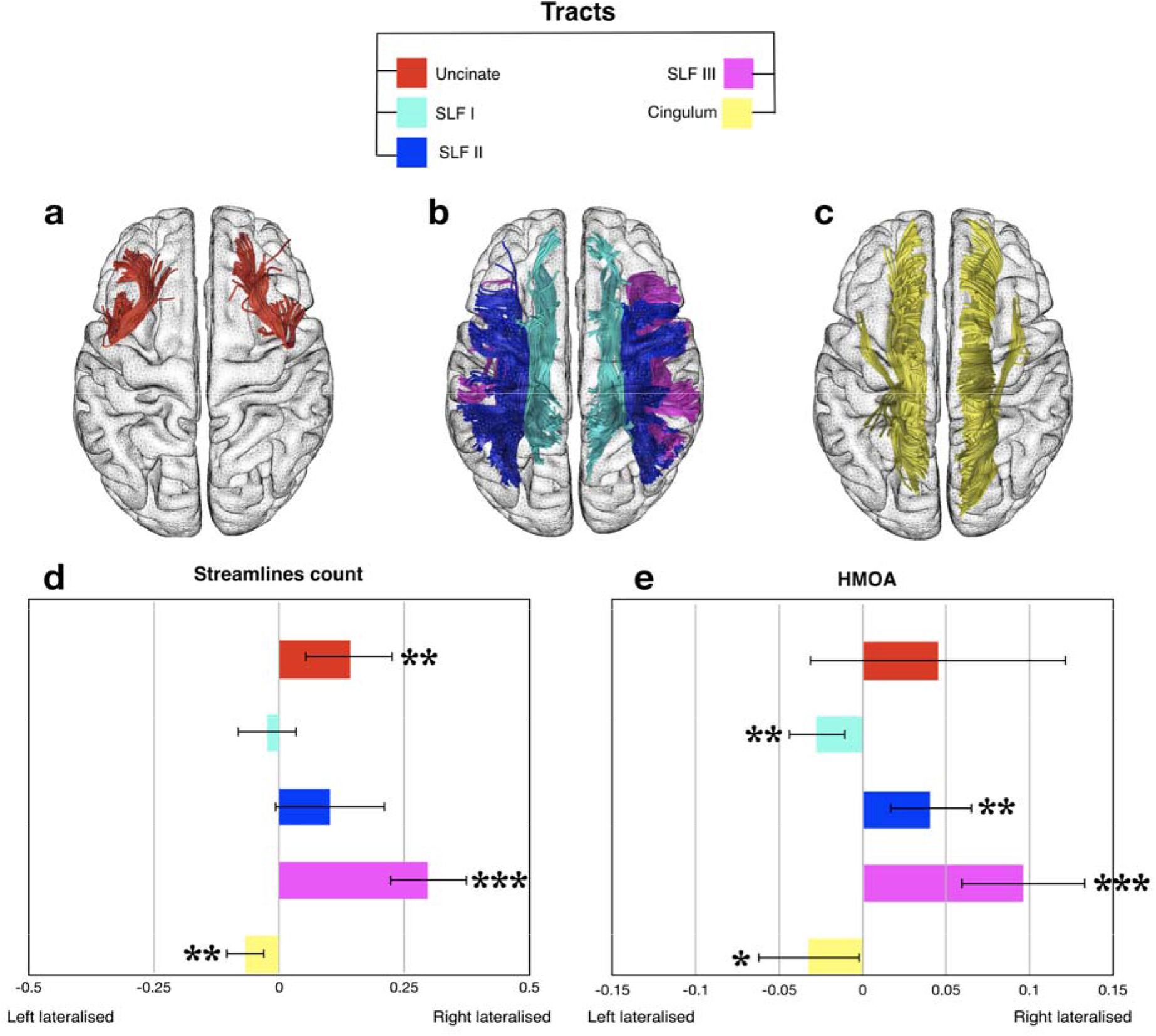
In vivo diffusion weighted imaging tractography dissection of a) uncinate fasciculus (red), b) superior longitudinal fasciculi, SLF I (cyan), II (blue), III (magenta) and c) cingulum fasciculus (yellow). d-e) Overall effect across the 33 participants for tracts asymmetries in terms of streamlines count (d) and HMOA (e). * p < 0.05, ** p < 0.01, *** p < 0.001. Note that the scales in d) and e) are different for visualization purposes.

We employed the software Trackvis (http://www.trackvis.org), which allows for visual inspection of the tracts and extracted a measure of their track count and hindrance modulated orientational anisotropy (HMOA)—a new index employed as a surrogate for tract microstructural organization (Dell’Acqua 2012). HMOA is preferable to fractional anisotropy (FA) in the case of algorithm solving fibre crossing. Indeed, HMOA is a true tract specific measurement, therefore HMOA values derived from dissections are not contaminated by other crossing tracts (i.e. partial volume effect).

For the isolation of the left and right uncinate fasciculi, a two ROIs approach was used. A coronal ROI was delineated around the white matter of the extreme capsule, together with an axial ‘AND’ ROI in the anterior temporal region.

A multiple region of interests (ROIs) approach was used to isolate the three branches of the superior longitudinal fasciculus (i.e. SLF I, SLF II and SLF III).

Three coronal ROIs were delineated around the white matter of the superior, middle and inferior/precentral frontal gyri, and another three ‘AND’ ROIs were delineated posteriorly in the parietal region.

Streamlines of the arcuate fasciculus projecting to the temporal lobe were excluded using an axial ‘NOT’ ROI in the temporal white matter (the arcuate is not part of the longitudinal system as it forms an arc to reach the temporal lobe)

Finally, a two ROIs approach was used to isolate the left and the right cingulum fasciculi. A sagittal ROI was delineated around the white matter of the cingulum for the left and the right hemispheres. Streamlines of the corpus callosum projecting to the hemisphere opposite to the ROI were excluded using a sagittal ‘NOT’ ROI in the corpus callosum.

ROI delineation illustrations are available as part of supplementary material of Rojkova et al. (2015).

An asymmetry index was calculated, based on the tract-averaged volumes (i.e. number of streamlines) and HMOA indices of the uncinate, SLF I, II and III and the cingulum using the following formulae

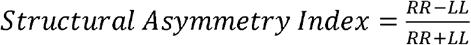

Here RR and LL represent the streamlines count and HMOA values for the tracts in the right (RR) and left (LL) hemispheres. A positive score indicated a larger tract in the right hemisphere whereas a negative score pointed to a leftward asymmetry.

### Statistics

First, behavioural and anatomical asymmetries were analysed independently using SPSS software (SPSS, Inc., Chicago, IL, United States of America) to perform one sample t-test against zero on the asymmetry indices of emotional expression and tract specific measurements.

Subsequently the relationship between asymmetry indices of emotional expression and structural asymmetry indices was determined using two hierarchical linear regression analyses (i.e. happy and sad conditions) with a maximum of 10 independent variables (i.e. asymmetry indices of streamlines and HMOA for uncinate, cingulum, SLF I, SLF II and SLF III) were introduced to a backward elimination analysis to identify significant regressors for predicting the asymmetry in the two conditions (i.e. happy or sad asymmetry indices.). This method first places all possible variables, as identified from the literature or driven by a specific hypothesis, in the model and calculates the contribution of each of them. The variable with the least contribution to the model is subsequently removed and statistical indices of the resulting model are re-estimated for the remaining variables. The contribution of the remaining variables is reassessed in an iterative way until the model reaches statistical significance.

## Results

### Behavioral measures

Replicability of the judges’ assessment was tested by splitting randomly the sample of 20 judges into two subsets of 10. The correlation between the average scores of the first 10 judges and the second 10 judges revealed a large effect of replication (Pearson correlation r = 0.75).

Subsequently, One sample t-test analysis applied to the full sample of judges (average score of n = 20) and posers (n = 33) did not reveal any difference between the left and the right side of the face in the happy (t(_32_)= 1.034; p = 0.309) and sad conditions (t(_32_)= 0.053; p = 0.958) (figure 2a) and there was no significant correlation between happy and sad conditions (r = 0.24; p = 0.171).

### Structural measures

*First, Paired sample t-test analysis demonstrated that there was no significant difference in ROIs size between the left and the right hemispheres (t_(32)_ < 1).*

One sample t-test analysis revealed a significant rightward hemispheric asymmetry for the number of streamlines of the uncinate fasciculus (t_(32)_= 3.24; p = 0.003) and the SLF III (t_(32)_= 7.84; p < 0.001), and for the HMOA of the SLF II (t_(32)_= 3.34; p = 0.002) and the SLF III (t_(32)_= 5.24; p < 0.001) (Figure 2d,e). A significant leftward hemispheric asymmetry was found for the cingulum in streamline counts (t_(32)_= −3.51; p = 0.001; Figure 2d) but not in HMOA after Bonferroni correction for multiple comparison (t_(32)_= −2.16; p = 0.039). A significant leftward hemispheric asymmetry was also found for the SLF I HMOA (t_(32)_= −3.27; p = 0.003; Figure 2e). P values were significant after Bonferroni correction for multiple comparisons for p < 0.005 (10 asymmetries indices).

**Figure 3:**
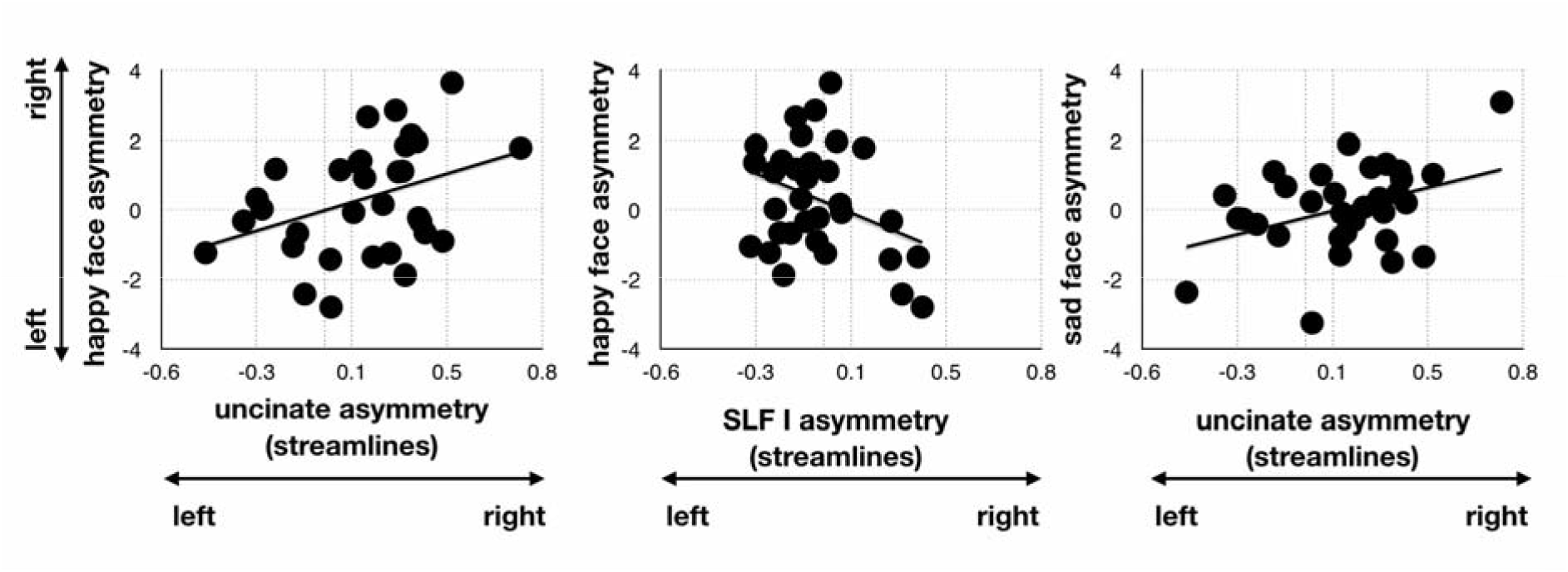
Dimensional relationship between measures of emotional expression asymmetries in the happy and sad condition and tracts asymmetries in terms of streamlines count.

### Behaviour-structure relationship

To confirm whether these anatomical asymmetries in white matter tracks significantly influence happy and sad asymmetry indices in our data set, a backward regression analysis was conducted. This method introduces all potential variables into the model at once before subsequently eliminating each independent variable, starting with the variable with the smallest partial correlation coefficient.

The analysis shows that when all variables are included, the model was not significant for predicting happy asymmetry indices [R^2^ = 0.538, F(10,22)=0.896, p = 0.552]. The following analyses removed step by step the asymmetry indices of the cingulum, SLF II and III track count and HMOA from the variables and the model became significant with the inclusion of uncinate and SLF I track count and SLF III HMOA [R^2^ = 0.527, F(3,29) = 3.727, p = 0.022]. Specifically, the asymmetry indices in uncinate and SLF1 track count appeared to have a significant contribution to the model (uncinate: B = 2.486, p = 0.019 and SLF I: B = – 3.516, p = 0.025) whereas the HMOA asymmetry index in SLF III did not appear to have an independent contribution (B = 1.013, p = 0.674).

Subsequently, the same analysis was carried on for predicting sad asymmetry indices and was not significant when all variables are included in the model [R^2^ =0.535, F(10,22)=0.882, p = 0.563]. The model became significant with the inclusion of asymmetry indices of the uncinate track count and SLF I HMOA [R^2^ = 0.468, F(2,30) = 4.208, p = 0.024]. Specifically, the asymmetry index in uncinate track count had a significant contribution to the model (B = 1.751, p = 0.036) whereas the HMOA asymmetry index in SLF I did not appear to have an independent contribution (B = 6.444, p = 0.128).

## Discussion

In this study we explored a classically reported effect of asymmetry in facial emotional expression and its relationship with the degree of lateralization of main white matter tracts. Three findings emerge from our work. First, the degree of asymmetry in emotional expression was not as prominent as reported in the literature, suggesting a significant inter-subject variability in the hemispheric prevalence for emotion expression. Second, the anatomical investigation was consistent with and clarified previous reports on the asymmetry of the uncinate, the SLF I, II and III and the cingulum. Third, we found significant relationships between the degrees of lateralization of main white matter tracts, with the asymmetry in the expression of happy and sad faces.

The degree of asymmetry in emotional expression we report is not as pronounced as reported in the literature on chimeric faces analyses (Sackheim et al, 1978). Compared to previous studies our analysis revealed a substantial variability among participants. In accordance with this, meta-analysis of lateralisation of emotional expression only report a weak effect at the group level (Skinner and Mullen, 1991). Some differences in our approach might explain this discrepancy. For instance, classical studies employing chimeric stimuli are based on a chosen subsample of a battery of already preselected emotional faces, which may have exaggerated the effect. Additionally, photographs were not emotional poses, but specifically requested facial movements (Ekman 1980). Here, chimeric stimuli were derived from undirected emotional poses of all the participants without any exception and might be more ecological and representative of the global population, revealing a highly variable level of posed emotions asymmetry across participants. Further, in previous approaches the chimeric faces were scored independently whereas in the current design we purposely put chimeric faces into competition to estimate the level of asymmetry in the expression at the individual level. Hence, our current paradigm is more appropriate to estimate asymmetry in the expression of emotion at the single subject level and therefore draw its relationship with asymmetries in the brain anatomy. Note that the side of initiation the emotion displays a stronger asymmetry than emotional expressivity on its own (Ross and Paulesu 2013). While initiation and intensity are different measures, we would like to stress that a combination of these two approaches might constitute the best behavioural marker to correlate with brain measures in future research.

Our results replicated preliminary evidence in the lateralization of previously described white matters tracts, providing further evidence on the distribution of these pathways in the healthy population. In line with previous reports, the uncinate fasciculus volume (i.e. streamline count) was significantly bigger in the right than in the left hemisphere (Hau et al. 2017, Highley 2002), the cingulum displayed a trend to leftward asymmetry (Thiebaut de Schotten et al. 2011b), while the SLF showed a dorsal to ventral right gradient of lateralization (Chechlacz 2015, Thiebaut de Schotten et al. 2011a).

Evaluating the relationship between the degree of lateralization of emotional expressivity with the asymmetries in various white matter bundles evinced partially overlapping neural correlates for happy and sad faces. The asymmetry of emotional expressivity was positively related to the volumetric asymmetry of the uncinate fasciculus. Albeit surprising, this ipsilateral effect might have been driven by the importance of the right uncinate fasciculus in emotional processing (Coad et al. 2017) and by the partially ipsilateral motor control of facial expression (Ross and Paulusu 2013). Being a key limbic tract, the role of this white matter tract in the domain of emotion is well established. As mentioned, this pathway connects the temporo-amygdala-orbitofrontal circuit, whose impairment leads to pathologies which are characterized by emotional and mood disturbances. Interestingly, the activity of the left amygdala has been reported to be stronger than its right hemisphere counterpart in response to emotional stimuli, in a series of studies that employed fearful and happy, but not sad, stimuli (Breiter et al, 1996). Additionally, recent evidences reported a mixed distribution of the lateralisation of the activation of regions during emotional processing, particularly left dominant for the ventral projections of the uncinate fasciculus (Karolis et al. 2019). The partial ipsilateral control of the face together with atypical lateralisation of the amygdala and the uncinate fasciculus in some emotional tasks suggests that the left amygdala might sometimes come in support of the right dominant emotional networks. The left amygdala occasional support would explain the ipsilateral relationship between the asymmetry of the uncinate fasciculus and the degree of asymmetry in emotional expression revealed in our study.

Contrary to the asymmetry of sad faces—which was only associated with the UF—the asymmetry in emotional expression of happy faces was inversely related to the lateralization of the SLF I, indicating a contralateral effect. Abnormalities in SLF microstructure have been reported as the most common white matter alteration in populations affected by emotion and mood disorders (Jenkins et al. 2016). Moreover, portions of the Supplementary Motor Area that are targeted by this white matter tract are involved in the production of orofacial movements, such as smiling, lip-licking and speaking (Kern et al. 2019). Taken together, these findings suggest a participation of the SLF I in the emotion domain, particularly in the expression of happiness.

This study is the first to the relevance of white matter asymmetries for the lateralization of emotional expressions in the left and right hemi-face. However, the pattern of results displayed both ipsilateral and contralateral effects, thus hinting at a more complex mechanism. The asymmetry in emotional expression was concordant with the lateralization of the uncinate so that a bigger left uncinate led to happier or sadder left hemiface (i.e. ipsilateral effect). Conversely, the lateralization of the SLF I was opposite to the asymmetry in happy expression so that a bigger left SLF1 led to a happier right hemiface (i.e. contralateral effect). While white matter asymmetries contribute to lateralized functions and behaviour (for review see Ocklenburg et al. 2014), recent trends in cerebral lateralization research propose a multi factorial, high dimensional perspective (Ocklenburg et al. 2014, Thiebaut de Schotten et al. 2019, Vingerhoets, 2019). Therefore, future research may address questions about the link between emotional expression and perception of emotions, as well as target brain asymmetries in grey matter and the potential influence commissural fibres.

## Limitations

We considered only happy and sad emotions in the present study. For practical reasons we only included these as they are universally shared, and thus provide more homogeneity across participants’ poses and are easily put in contrast. Also, several reports indicate that happy faces are usually more distinctive from neutral faces than sad faces (smiling causes an expansion of the face), and have a perceptual advantage in detection and recognition (Becker et al. 2011, Hodsoll et al. 2011).

Additionally, the expressed emotions were emotional poses rather than spontaneous. As this is a highly criticized issue (Ekman 1980), we would like to clarify that our participants were requested to express either happiness or sadness, without giving them indications on how to do so. We chose emotional poses in this study as there is support for the notion that voluntary control of facial movement is important for the intensity of spontaneous facial expressions (Berembaum and Rotter 1992), and because full-frontal face view photographs are crucial for the study of emotional expression. Furthermore, because stimuli were only presented in competition, we could not directly assess the absolute intensity of emotion expressed on each chimeric face and therefore limited our analysis to asymmetries.

Another point concerns the small number of participants who took part in the present study. Whilst a group of 15 is usually accepted in the field as a sufficient sample size to produce group results, increasing sample size might improve the likeliness to reveal small effects as significant. Here, power calculation based on our data indicated that one would need a minimum sample of 337 posers for the happy condition and 11000 posers for the sad condition in order to obtain a significant difference. The purpose of our study was, however, focused on variability (Thiebaut de Schotten and Shallice 2017) and revealed that left hemiface dominance for emotional expression is true in some but not all participants and that this variability correlate with anatomical asymmetries. Thus, increasing our sample size would have probably only inflated the significance of our findings.

## Conclusion

The present work sought to explore the relationship between the hemispherical dominance in the posed expression of happy and sad emotion with the asymmetry of white matter tracts of the human brain. We revealed an unexpected large inter-subject variability in the asymmetrical expression of emotion, which was related to the lateralization of the Uncinate Fasciculus and the SLF I and brought novel speculation on the emotional contributions of these white matter tracts. Future studies could expand the knowledge on the nature of the interaction between brain structural connectivity and emotions with keen attention to the implications for clinical populations affected by mood disorders

## Acknowledgments

We thank ERASMUS for supporting SI. This project has received funding from the European Research Council (ERC) under the European Union’s Horizon 2020 research and innovation programme (grant agreement No. 818521). The research leading to these results also received funding from the “Agence Nationale de la Recherche” [grants number ANR-13-JSV4-0001-01] and from the Fondation pour la Recherche Médicale (FRM). Additional financial support comes from the program “Investissements d’avenir” ANR-10-IAIHU-06.

